# Non-heritable variation in individual fitness adds stability to neutral theories in ecology and evolution

**DOI:** 10.1101/379776

**Authors:** M. Gabriela M. Gomes, Jessica G. King, Ana Nunes, Nick Colegrave, Ary A. Hoffmann

## Abstract

Neutral theories in ecology^1^ and evolution^2^ contend that high diversity of natural communities and high rates of molecular evolution conform to models where individuals have equal fitness and mutations have no effects. Demographic stochasticity makes community and population compositions inherently unstable under these models, with overall levels of diversity being maintained by random processes. This is in contrast with niche and adaptive theories which emphasize differences between species or genotypes as the key to their coexistence^3^. Here we show that non-heritable variation in individual fitness within species or genotypes can stabilize coexistence without evoking niche differentiation. We construct two classes of mathematical models based on experimental evidence: (1) bacterial growth with variation in cell longevity^4,5^; and (2) microbial transmission in a host population with variation in host susceptibility^6–11^. We find stable coexistence of 2 bacterial species in the first model under a single oscillating resource, and 3 or more in the second with independent distributions of host susceptibility to the various microbial species. We discuss the implications of these findings for the interpretation of common measures of relative fitness and for the maintenance of biodiversity.

---

Growing populations of genetically identical bacteria placed under selective antibiotic pressure exhibit a decline over time in their rates of mortality^4,5^. When observed in time frames that are too short to reflect inherited mutations, this pattern has been attributed to variation in non-inherited sensitivity of individual cells to the antibiotic. This may be induced by variation in rates of cell division, especially for antibiotics that reduce the viability of new cells.

We modify established mathematical formalisms representing bacterial population dynamics^12–14^ to include individual variation in rates of cell division. Different arguments will be made about mother and daughter cells, and therefore the process needs to be separated into cell death and birth in our models. In the simplest instance of unlimited growth, this is written as

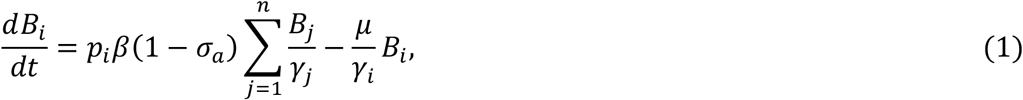

where *B_i_*, for *i* = 1, …, *n*, denote the concentration of bacteria with longevity factor *γ_i_*, in a fraction *p_i_* of all births, purporting a distribution with mean 〈*γ*〉 = Σ*_i_p_i_γ_i_* = 1, variance 〈(*γ* − 1)^2^〉 = Σ_i_P_i_(y_i_ – 1)^2^, and coefficient of variation 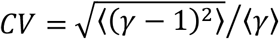 treated as a varying parameter. To ensure that replication is by binary fission we impose *β* = 2*μ*, and to enable fitness comparisons across distributions we denote the mean rate of cell division by *M* = 〈*μ*/*γ*〉. Antibiotic pressure is represented as a reduction in the viability of newborn cells (1 − *σ_a_*). We treat model (1) as a building block which can be coupled to various models of resource provision as desired (Methods).

Balaban *et al.* (2004) investigated the persistence of inherently sensitive cells when a population of genetically identical bacteria is exposed to an antibiotic stress, a phenomenon first observed in the early days of penicillin use^15^. The authors described mathematically the dynamics of surviving cells by switching mechanisms between a majority of rapidly growing “normal” cells and a minority of slowly growing “persister” cells. Coupling model (1) to a system of continuous resource provision (Methods, model (4)-(5)) we reproduce the same results without evoking phenotypic switches (Fig. 1). The model is solved numerically without antibiotic (*σ_a_* = 0) through the exponential phase until a stationary state is established, at which stage the antibiotic is introduced (*σ_a_* = 0.9). In the absence of variation in cell longevity, the antibiotic causes an exponential decay in cell density (solid black line in Fig. 1a). The slightest variation in longevity induces a form of cohort selection that results in decelerated population decay (illustrated by the dashed black line in Fig. 1a generated with a coefficient of variation of 0.05). The greater the variation, the greater the deceleration (magenta line in Fig. 1a generated with the coefficient of variation set to 3). Fig. 1b shows the action of cohort selection. As time under antibiotic increases, the faster dividing cells become rarer in the population (i.e. the fraction of persisters increases). The original distribution of cell longevity is continuously being reset through new births, but viability is generally low due to antibiotic pressure leading to an accumulation of long-lived cells. The same phenomenon occurs regardless of whether the population is structured into two discrete groups or shows a continuous distribution of longevity factors (Fig. 1d, e). Indeed, different types of survival curves, as reported by Balaban *et al.* (2004), can be obtained by concordantly setting the distribution of longevity factors without needing additional switches or other processes^16–18^. These ideas apply to growing cell populations more generally and may be extended to describe failure of treatments in cancer patients^19^, as well as a wide variety of bet-hedging strategies in nature^20^.

**Fig. 1:**
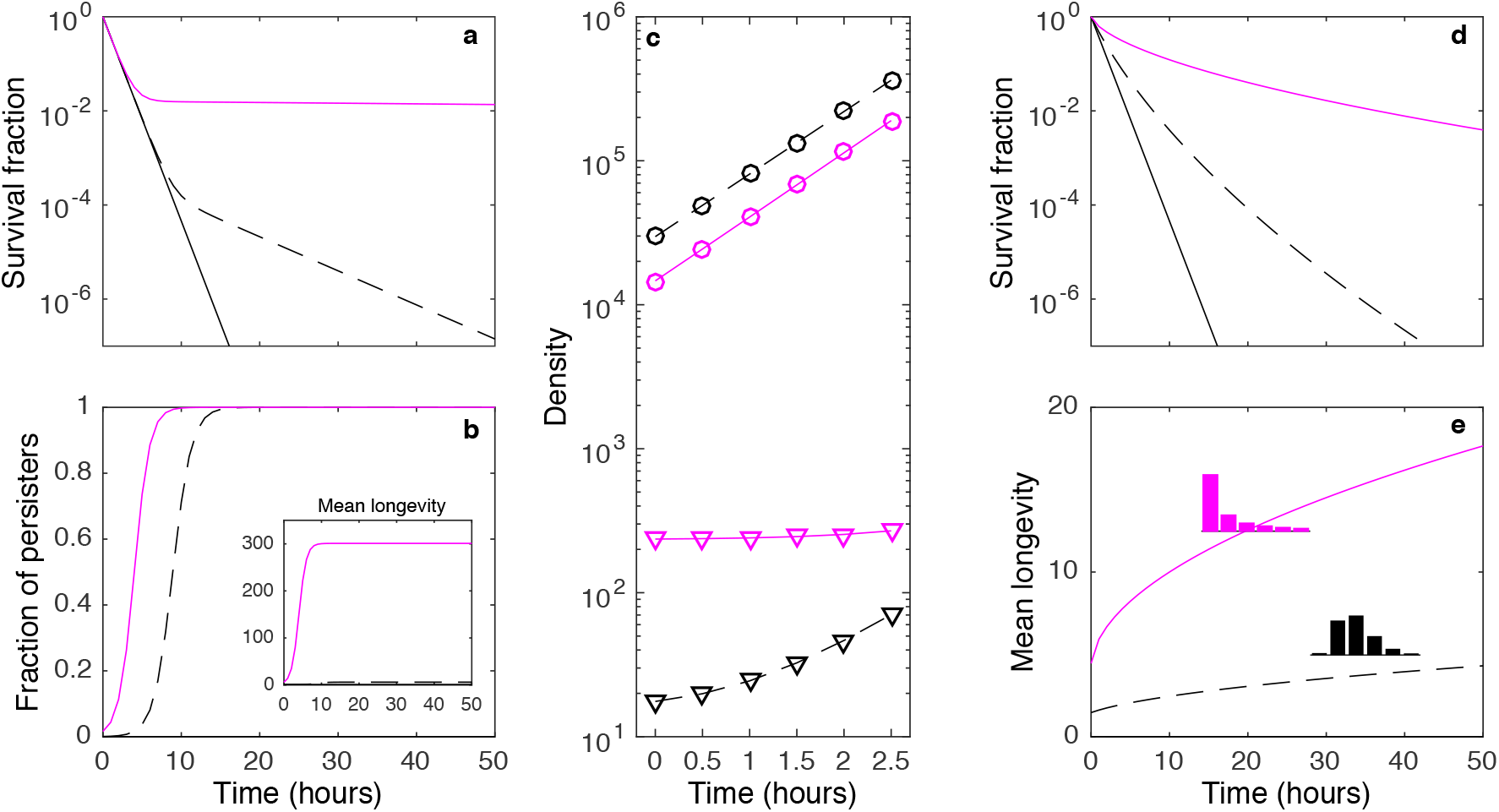
Bacterial persistence to antibiotic treatments. **a**, **b**, **c**, Solutions of model (4)-(5) with two-group distributed longevity factors, *γ*. The fraction of cell births entering the high-longevity group was set to 0.0001. Two distinct coefficients of variation are represented: *CV* = 0.05 (dashed black), and *CV* = 3 (magenta). The solid black curve represents a homogeneous population: *CV* = 0. A pre-antibiotic phase (*σ_a_* = 0) was simulated with *c* = 2 and *ρ* = 0.003, until a stationary phase was established. Stationary phase solutions were used to simulate: **a**, **b**, antibiotic introduction by setting *σ_a_* = 0.9 and turning off the chemostat flow (*ρ* = 0); and **c**, growth without antibiotic by keeping *σ_a_* = 0 and setting *R*(0) = 10^6^. Curves punctuated by circles represent total populations, whereas triangles refer to persistent fractions. **d**, **e**, Solutions of model (4)-(5) with gamma distributed longevity factors and two distinct coefficients of variation: *CV* = 0.5 (dashed black), and *CV* = 2 (magenta). Other parameters: *M* = 1, *ϕ*(*R*) = *R*/1 + *R*).

Fundamental to the results above is a notion of genotype fitness that is wider than that commonly used. By accommodating explicitly for individual variation in a fitness trait, the *effective* genotype fitness is not fixed; it is determined by how the *intrinsic* fitness distribution is molded by stress. Here we consider environmental stresses which reduce cell viability at birth (Fig. 2a, b), reduce survival at any age (Fig 2c, d), or affect cell division rates (Fig 2e, f). Considering mutants derived from ancestral genotypes under multiple scenarios, we describe the possible patterns which may occur when fitness ratios are measured (*r_m_*/*r_a_*, where *r_a_* and *r_m_* denote ancestor and mutant growth rates, respectively, measured at some time point during exponential phase). First, we assume that the phenotypic variance of the ancestor is negligible compared to the mutant and find this to result in effective fitness ratios (solid colored lines in Fig. 2a, c, e) that are consistently lower than the ratios of the respective intrinsic fitness (dashed lines) and that decrease with stress. This trend is common^21^, but the reverse has also been observed^22^ and occurs in our framework when mutants are less variable than their ancestors (Fig. 2b, d, f). Genetic stresses can induce similar phenomena and affect measurements of epistasis^23^. Any mutation with an effect on fitness sets a differential in stress levels between ancestral and mutant genotypes, introducing a bias in the assessment of the intrinsic effects of additional mutations. If an initial mutation increases fitness variance, for instance, a second mutation may appear to have a smaller effect without evoking epistasis between the mutations. These trends must impact the estimation of distributions of fitness effects of mutations^24^.

**Fig. 2:**
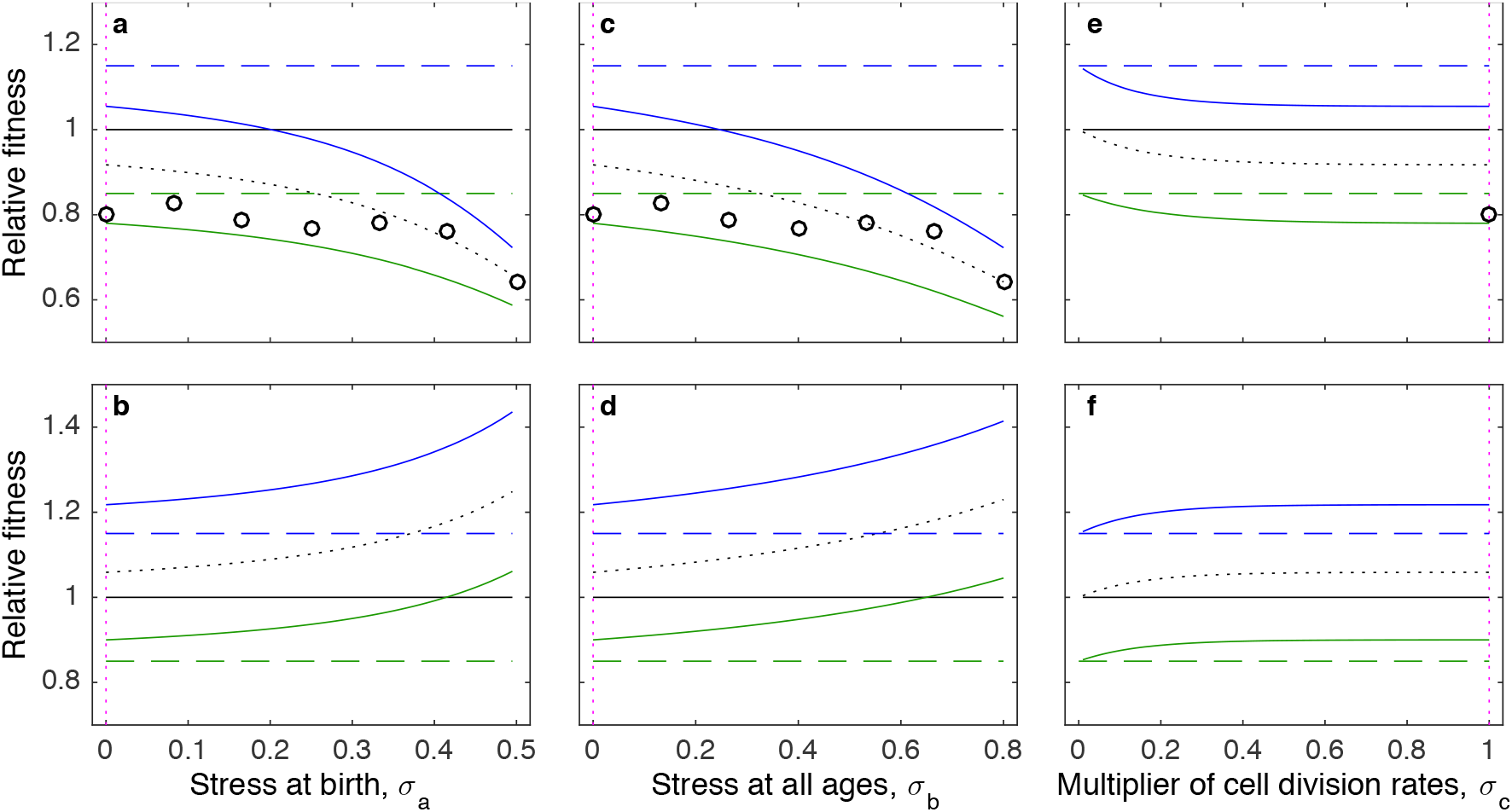
Relative fitness across stress gradients. Relative growth rates between mutant and ancestral genotypes calculated from model (1) at time *t* = 6*h* with: (**a**, **c**, **e**) *M* = 1 and *CV* = 0 (black, ancestral genotype), *M* = 0.85 and *CV* = 3 (green, mutant), *M* = 1.15 and *CV* = 3 (blue, mutant), and *M* = 1 and *CV* = 3 (black, dotted); (**c**, **d**, **f**) *M* = 1 and *CV* = 1 (black), *M* = 0.85 and *CV* = 0 (green), *M* = 1.15 and *CV* = 0 (blue), and *M* = 1 and *CV* = 0 (black, dotted). Stress was implemented in three ways: (**a, b**) reduction in cell viability at birth (parameter *σ_a_* in model (1)); (**c, d**) increase in cell mortality at all ages (parameter *σ_b_* in model (6); Methods); or (**e, f**) factor affecting the rate of cell division (parameter *σ_c_* in model (7); Methods). Vertical dotted lines (magenta) indicate where the three axes (σ_a_, *σ*_b_, *σ*_c_) intersect. The fraction of cell births entering high-longevity groups is set to 0.09. Dots indicate relative fitness measurement on linear stress gradients as in Kraemer *et al*.(2016).

In light of these findings, we argue that when variation in individual fitness exists, the intrinsic fitness ratio between genotypes and corresponding selection coefficient (1 − *r_m_*/*r_a_*) cannot be measured directly but can be estimated by fitting a curve to effective fitness ratio measurements across stress gradients (as recently proposed for relative disease risks and vaccine efficacy^25^). This is because effective and intrinsic fitness ratio curves only coincide in a limit where stress is zero, which here signifies immortality and thus no growth rates to be measured.

Current definitions of fitness and associated measures in population biology are too narrow to accommodate the phenomena described above. This is strikingly conveyed by Fig. 2 where fitness curves of two genotypes measured across a stress gradient effectively cross at some critical stress value (solid blue and black curves in Fig. 2a, c, and green and black curves in Fig. 2b, d) where the selection coefficient is zero. The populations differ, however, in intrinsic fitness and the crossing is due to the action of cohort selection. Unaccounted phenotypic variation within genotypes is therefore capable of stabilizing coexistence of multiple genotypes and unexpectedly affect patterns of genetic variation^26^, especially when levels of stress fluctuate. Firstly, genetic diversity of fitness traits could be higher than expected even though these traits typically have low heritability^27,28^. Secondly, genetic drift and other random processes involving population extinctions are likely to be slower than predicted by current models (similar to what has been noted in the context of species conservation^29^).

We now turn to a classic debate in community ecology about whether the high diversity of species that exists even when competing for the same resources is attributed to “equalizing” (neutral theory) or “stabilizing” (niche theory) mechanisms^3^. The neutral theory^1^ posits that individuals, irrespective of species, are basically identical in their fitness and their interactions, and community dynamics are driven by demographic stochasticity and speciation^1^. The niche theory, by contrast, proposes that species differ in their niches^30,31^ and that the negative effects of intraspecific individual interactions are larger than those due to interspecific interactions. This dichotomy can also be presented as a contention between stochasticity and determinism^32^. Our results above, however, suggest that introducing individual variation into essentially neutral models can lead to coexistence that is deterministically sustained without stabilizing mechanisms.

Model (8)-(11) extends the models used above by accommodating three species and introduces an oscillating concentration of the single resource entering the system. Fig. 3 shows various tongue-shaped regions (in yellow) outlining stable coexistence between pairs of species. Previous studies have described coexistence in similar systems^12–14^, but relied on different species having different viability functions (*ϕ*(*R*), where *R* is the concentration of resources and *ϕ*(*R*) the viability of newly born cells), thus complying with the niche theory. In contrast, the mechanism we describe here relies on cohort selection acting on individual variation in longevity under oscillating resources, and is neutral when framed within current theories, which are essentially blind to intraspecific individual variation. When resources are low, high-longevity cells are at an advantage and so are species exhibiting higher variation, whereas under abundant resources species with less variation have an advantage because they effectively grow faster, generating a pattern that can be interpreted as negative frequency-dependent selection^33^. This mechanism does not appear to sustain coexistence of more than two species, but fitness is typically governed by many traits and variation in any of them can open possibilities for coexistence.

**Fig. 3:**
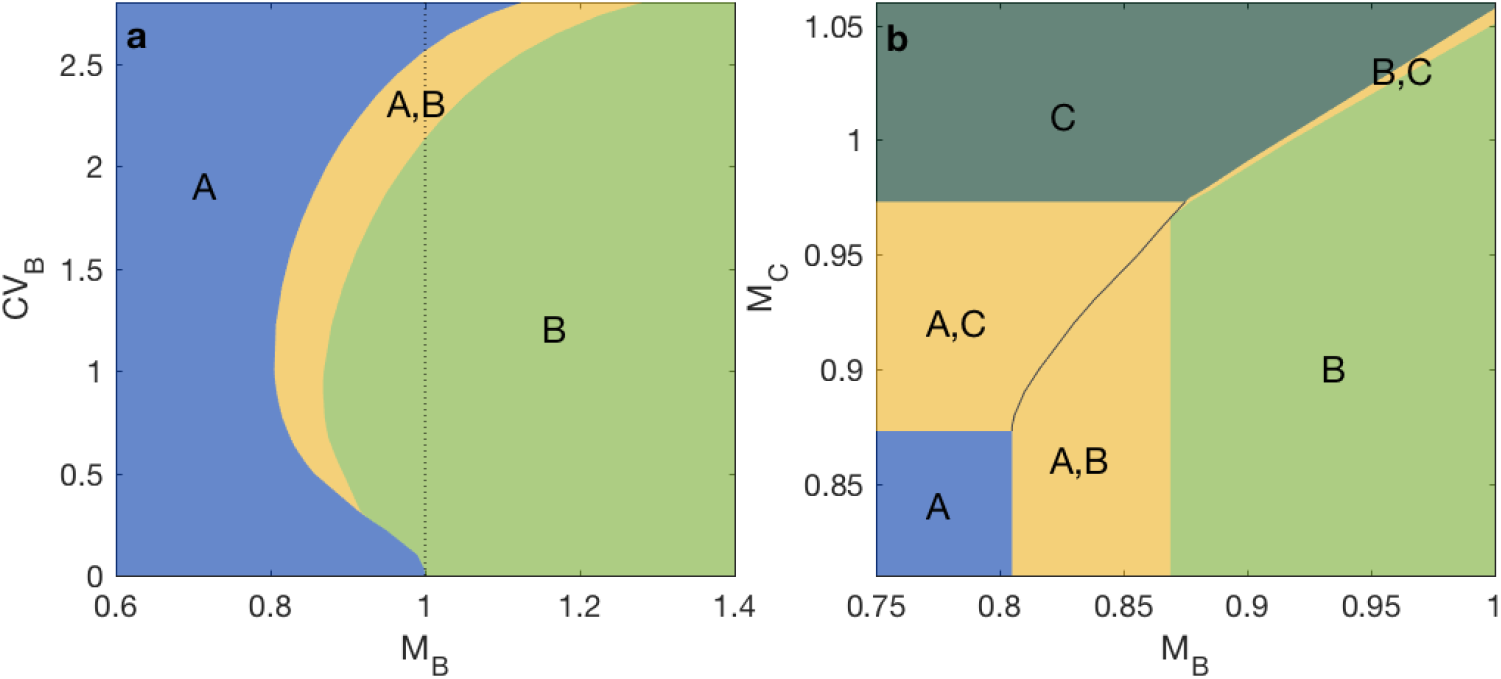
Stable coexistence of microbial species in an oscillating chemostat. Model (8)-(11) was solved numerically with two (**a**) and three (**b**) species (Methods). Yellow tongues represent regions of stable coexistence among the indicated species. All species have the same cell viability function *ϕ*(*R*) = *R*/(1 + *R*), the chemostat flow is set to *ρ* = 0.1, and the concentration of resources in the input flow oscillates as *c*(*t*) = 3[1 + cos(2*πt*/24)]. Other parameters: (a, b) *CV_A_* = 0 and *M_A_* = 1; (b) *CV_B_* = 1 and *CV_C_* = 2.

Shifting from longevity to resource accessibility, for example, and its effect on birth viability, we construct a model for the transmission of microbial species (or strains) in a host population. For a single species, variation in resource accessibility is assumed to arise from variation in susceptibility of individual hosts to microbial colonization

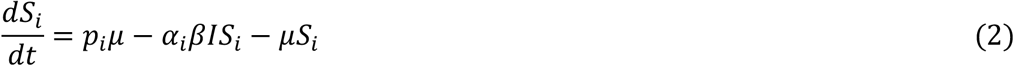

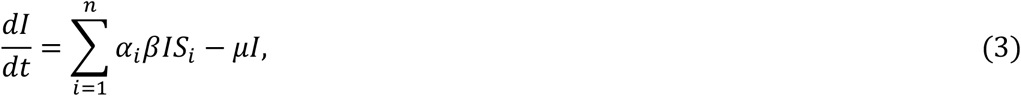

where *μ* is the host death and birth rate, *β* is the effective contact rate between infected and susceptible hosts, *α_i_* is the susceptibility factor of hosts *S_i_* that enter the system as a fraction pi of all births, purporting a distribution with mean 〈*α*〉 = Σ_*i*_*p_i_α_i_* = 1, variance 〈(α− 1)^2^〉 = Σ_*i*_*p_i_*(*α_i_* − 1)^2^, and coefficient of variation 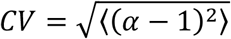 treated as a varying parameter. Systems such as this may provide more accurate descriptions of infectious disease dynamics than their homogeneous analogues^7,10,11^. Based on this building block we construct a family of high-dimensional models for the circulation of *N* microbial species, each independently facing a host population with some susceptibility distribution (Methods). Coexistence regions for 2 and 3 species are shown in Fig. 4 and can be generated inductively for any natural number *N*.

**Fig. 4:**
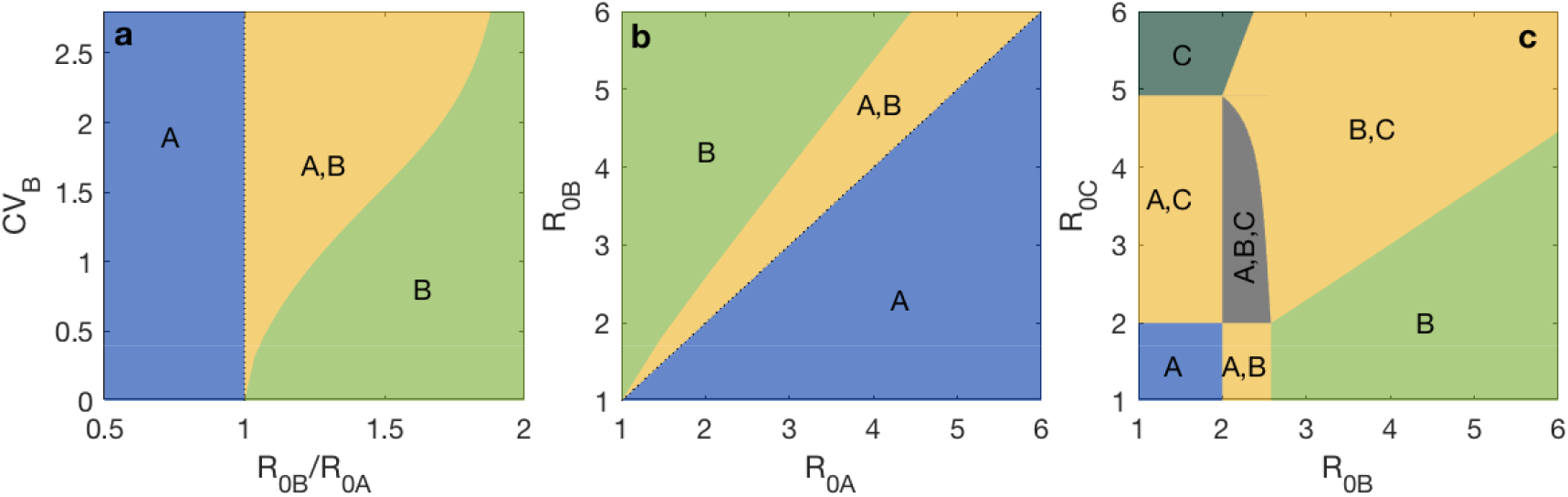
Stable coexistence of microbial species colonizing a host population. Model (12)-( 15) was solved analytically with two (**a, b**) and three (**c**) species (Methods). Yellow regions represent conditions for 2-species stable coexistence as indicated, while 3-species coexistence is found in the gray zone (c). Other parameters: (a, b, c) *CV_A_* = 0; (b, c) *CV_B_* =1; (c) *CV_C_* = 2 and *R*_0*A*_ = 2.

Until now stabilizing mechanisms that sustain coexistence have been tied to species as homogeneous static entities^3,34^. We challenge this idea by showing how measured variation in individual fitness can stabilize neutral coexistence across ranges of environmental conditions and set a scenario where the debate between neutral *vs* niche theories of biodiversity becomes immaterial. In parallel, we also find that non-heritable phenotypic variation within genotypes can stabilize coexistence of multiple genotypes thereby sustaining substantial heritable variation despite being under stabilizing selection^26^.

According to neutral theories of diversity at genetic^2^ and species^1^ levels, the heritable variation that continually arises through mutation and migration is subject to stochastic processes that allow transient and therefore unstable coexistence of multiple genotypes or species. Stabilization of coexistence, on the other hand, can arise from specialization of genotypes and species in separate fitness peaks and ecological niches, respectively. Throughout this manuscript, we have argued towards a unifying theory of biodiversity, at genetic and species levels, that is both neutral and stable and that is described by deterministic processes. This is achieved by relaxing a previously implicit assumption that individuals within a genotype or species are phenotypically identical. Non-heritable variation in fitness triggers the action of cohort selection providing genotypes or species with a buffer that decreases or even hinders the effects of selection between them and promotes biodiversity.

## METHODS

### Bacterial growth models

Limited bacterial growth is induced by limiting resources. For a bacterial genotype (or species) maintained in continuous culture in a chemostat, the model is written as

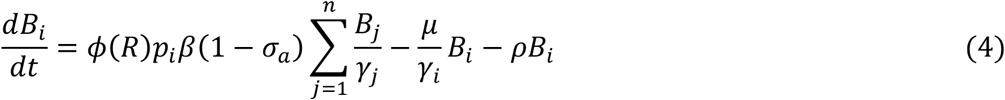

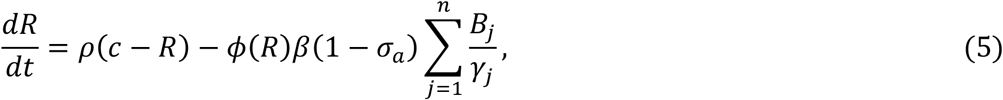

where *R* is the concentration of resources in the chemostat, *c* its concentration in the input flow, *ρ* the rate at which medium enters and leaves the chemostat, and *ϕ*(*R*) is a nonnegative increasing function between 0 and 1 describing the viability of newly born cells as a function of resource availability, here assumed a Hill function *ϕ*(*R*) = *R*/( 1 + *R*). The remaining parameters and variables are as in model (1). This model was used to generate Fig. 1. Numerical solutions of system (4)-(5) were obtained for populations structured into either two longevity groups (a majority group with index 1 and a smaller group with greater longevity, indexed by 2, to account for persistent cells) or continuous gamma distributions discretized into hundreds of longevity groups.

Antibiotic treatments were implemented as stresses that reduce the viability of newly born cells by a factor *σ_a_* (model (1)). Another form of environmental stress would be a mortality rate factor affecting all cells irrespective of age:

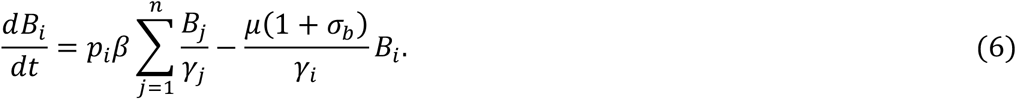

More favorable environments, by contrast, may be argued to reduce the need for cells to divide and thereby slow down the rate of cell division:

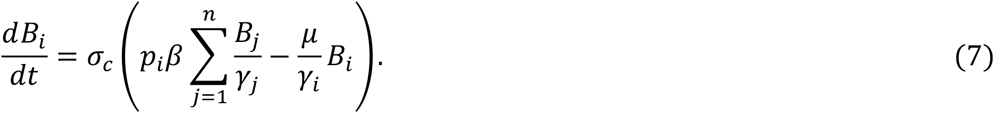

The three models ((1), (6) and (7)) were used to assess the sensitivity of fitness measures to selection on non-heritable variation.

To illustrate how distributions of longevity affect competition outcomes, we added two more species to the basic model without stress parameters (assuming one species homogeneous without losing the generality of possible outcomes), and set the concentration of resources in the input flow to oscillate with a 24*h* period, *c*(*t*) = *c*_0_[1 + cos(2*πt*/24)]. The resulting model used to generate Fig. 3 is written as

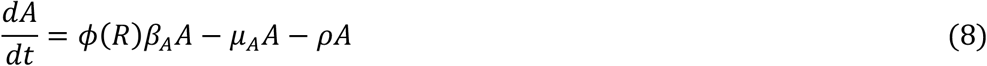

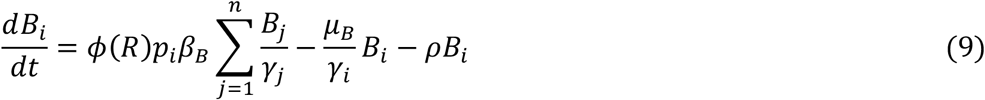

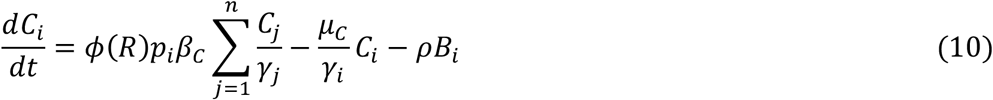

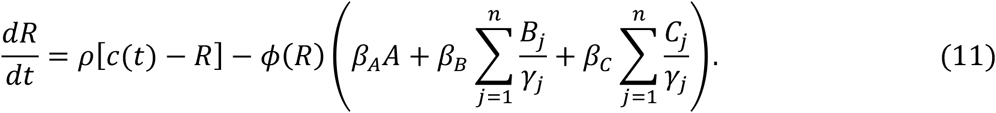

In this case, numerical solutions were obtained for populations structured into two longevity groups only. The formalism, however, is generic and can accommodate other discrete or continuous distributions.

### Host population models

The model for three microbial species (*A, B* and *C*) circulating in a host population, presenting *n_A_*, *n_B_* and *n_c_* susceptibility groups, respectively, to each species is written as

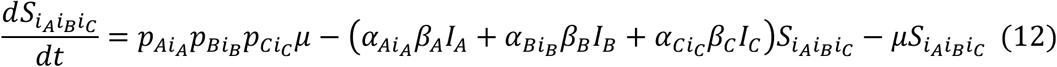

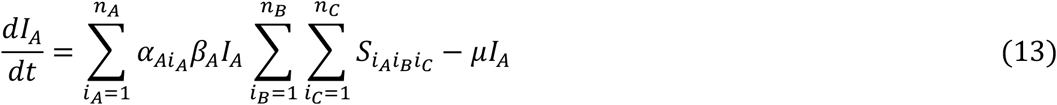

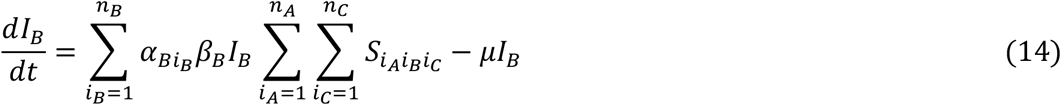

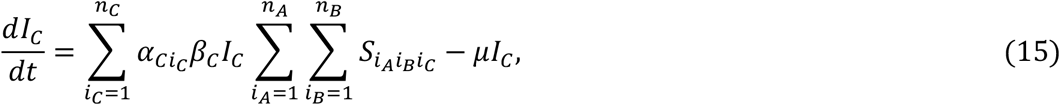

where *μ* is the host death and birth rate, *β_A_*, *β_B_* and *β_C_* are the effective contact rates between hosts infected by species *A, B* and *C* and susceptible hosts, *α_Ai_A__*, *α_Bi_B__* and *α_Ci_C__*, for *i_A_* = 1,…, *n_A_*, *i_B_* = 1, …, *n_B_* and *i_C_* = 1, …, *n_C_* are the susceptibility factors of hosts *S_i_A..__*, *S_.i_B·__* and *S_..i_C__*, who enter the system as fractions *p_Ai_A__*, *p_Bi_B__* and *p_Ci_C__* of all births, purporting distributions with mean 〈*α_A_*〉 = Σ*_i_A__p_Ai_A__α_Ai_A__* = 1, 〈*α_B_*〉 = Σ*_i_B__p_Bi_B__α_Bi_B__* = 1 and 〈*α_C_*〉 = Σ*_iC_C__p_Ci_C__α_Ci_C__* = 1, variance 〈(*α_j_* − 1)^2^〉 = Σ*_i_A__p_Ai_A__*(*α_Ai_A__* − 1)^2^, 〈*α_B_* − 1)^2^〉 = Σ*_i_B__p_Bi_B__*(*α_Bi_B__* − 1)^2^ and 〈*α_C_* − 1)^2^〉 = Σ*_i_C__p_Ci_C__*(*α_Ci_C__* − 1)^2^, and coefficients of variation 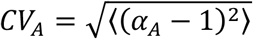, 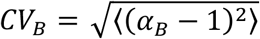 and 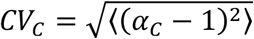 treated as varying parameters. The species-specific basic reproduction numbers are *R*_0*A*_ = *β_A_*/*μ*, *R*_0*B*_ = *β_B_*/*μ* and *R*_0*C*_ = *β_C_*/*μ*.

In the special case where the host population is homogeneously susceptible to *A* (*n_A_* = 1), heterogeneous to *B* with two susceptibility groups (*n_B_* = 2), and *C* is absent (*n_C_* = 0), coexistence occurs when *R*_0*A*_, *R*_0*B*_ > 1 and 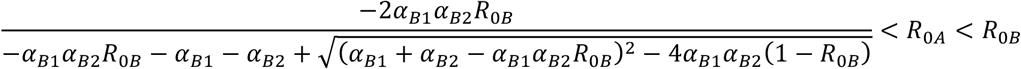.

These conditions were used to delineate the 2-species coexistence tongues in Fig. 4a, b. In the case of 3 species (*n_C_* = 2), the conditions for coexistence among the various pairs are analogous and were used to partially generate Fig. 4c. To complete the figure, the 3-species coexistence region, which exists for *R*_0*A*_, *R*_0*B*_, *R*_0*C*_ > 1, is bounded by the straight lines *R_0B_= R_0A_* and *R*_0*C*_ = *R*_0*A*_ (where species *B* and *C*, respectively, become absent), and by a third line, where species *A* becomes absent. This line can be obtained analytically by assuming 3-species coexistence and then setting the equilibrium abundance of species *A* equal to zero.

The extension of the model to *N* species is straightforward although the notation becomes dense:

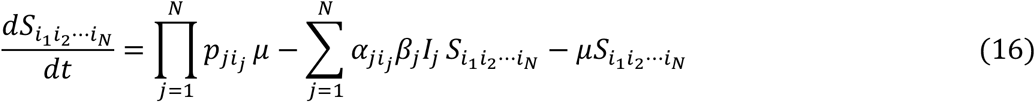

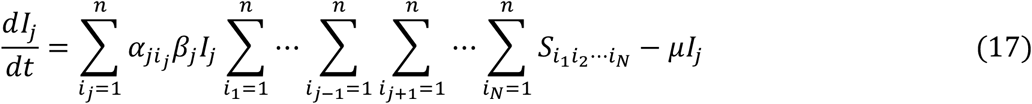

where *μ* is the host death and birth rate, *β_j_*, for *j* = 1,…, *N* is the effective contact rate between hosts infected by species *j* and susceptible hosts, *α_ji_j__*, for *i_j_* = 1,…, *n_j_*, are the susceptibility factors of hosts, *S_…i_j⋯__*, who enter the system as fractions *p_ji_j__* of all births, purporting distributions with mean 〈*α_j_*〉 = Σ*_i_j__p_ji_j__α_ji_j__* = 1, variance 〈(α_j_ − 1)^2^〉 = Σ*_i_p_ji_j__* (*α_ji_j__* − 1)^2^, and coefficients of variation 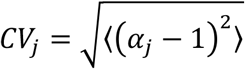 treated as varying parameters. The species-specific basic reproduction numbers are *R*_0*j*_ = *β_j_*/*μ*. In the special case where the host population is homogeneously susceptible to species 1, we find an *N*-species coexistence region with all *R*_0*j*_, > 1. This region has a simple geometry in the *R*_0*j*_, space that generalizes the 3-species coexistence. It is bounded by the hyperplanes *R*_0*j*_= *R*_0*A*_, for *j* = 2,…, *N*, and by a hypersurface that can be obtained as before by setting to zero the coexistence abundance of species 1. This coexistence region persists when we allow for heterogeneous susceptibility to species 1 as well.

## Acknowledgements

MGMG received funding from Fundação para a Ciência e a Tecnologia (IF/01346/2014) and AAH received fellowship funding from the National Health and Medical Research Council.

## Author contributions

M.G.M.G. designed the study and drafted the manuscript; all authors wrote the paper.

## Competing interests

The authors declare no competing interests.

## Data availability

Not applicable.

## Supplementary Information

Not applicable.

